# Effector selection precedes movement specification: evidence from repetition effects in motor planning

**DOI:** 10.1101/2024.10.23.619895

**Authors:** Christian Seegelke, Tobias Heed

**Affiliations:** Department of Psychology, University of Salzburg, Salzburg, Austria; Centre for Cognitive Neuroscience, University of Salzburg, Salzburg, Austria

**Keywords:** motor planning, repetition effects, action selection, effector specificity, limb choice

## Abstract

Motor performance is influenced by movements that were performed shortly prior. For example, reaction times (RTs) for successive movements are reduced when executed with the same effector, even if the specifics of the consecutive movements differ. These findings have been taken to support the notion that repetition effects in motor planning reflect the involvement of effector-specific motor plans. However, previous studies have confounded motor and visual aspects of repetition: movements have typically been instructed via visual cues, and movement repetition, therefore, implied repeating also the visual cue, so that the latter may be (at least partly) responsible for the observed RT effects. In the present study, participants performed two movements in succession, a prime and a probe action, either with their left or right hand and in one of two directions, inward or outward relative to the body midline. We used different cues for prime and probe actions, so that movement repetitions did not involve repetition of the visual cue. Participants initiated successive same-limb movements faster than different-limb movements, but this RT advantage was smaller than observed in previous work. Moreover, repeating movement direction also led to a decrease in RT, though only in combination with hand repetition. Whereas these findings imply that visual cue repetition can contribute to accelerated RTs in movement repetition, they confirm that the recent motor history affects motor planning. Furthermore, they support the idea of a hierarchical framework of motor planning in which effector selection precedes specification of motor parameters.

## Introduction

In daily life, we constantly make decisions about possible courses of actions, such as which hand to use for a given task. For instance, you may open a door with one or the other hand, depending on whether one hand is carrying a bag. Decisional processes related to effector selection are typically investigated via paradigms in which participants must choose a hand (left, right) with which to attain a motor goal such as reaching to a spatial location. Numerous factors can bias this choice, such as participants’ handedness (Bryden et al., 2000), future task demands (Coelho et al., 2014), effort (Schweighofer et al., 2015), and the recent motor history (Oliveira et al., 2010; Schütz & Schack, 2020; Valyear et al., 2019). The influence of motor history is apparent as a tendency to use the same hand for consecutive actions. The most common explanation for such motor history effects is that they reflect recycling of previously generated motor plans such that parameters of previously generated plans are reused for current actions (Rosenbaum et al., 2007; Schütz & Schack, 2013).

In line with this proposal, reaction times (RTs) are faster for the second of two consecutive actions – hereafter referred to as prime (i.e., the first) and probe (i.e., the second) action –, when the two actions are performed with the same effector. This finding holds even when the two actions involve different movements; thus, the RT improvement reflects a limb repetition effect. In contrast, repeating the same movement, but with a different effector, did not yield shorter RTs (Seegelke et al., 2021; Valyear & Frey, 2014), suggesting that limb repetition effects reflect effector-specific coding.

On a neural level, these effector-specific RT advantages are thought to result from persistent changes within neural populations that are dedicated to a given effector. Residual activity from previous movements would thus require less additional activity for initiation of a movement with the same limb, hence yielding shorter RTs in effector repetition trials. This idea is supported by the finding that decreased movement latencies in effector repetition trials were accompanied by reduced fMRI activity in bilateral posterior parietal cortex (Valyear & Frey, 2015). Furthermore, computational modelling based on the framework of drift diffusion suggests that the RT advantage for limb reuse are due to a shift of the starting point of evidence accumulation (Seegelke et al., 2021), which lends further support to the idea that repetition effects reflect persistent changes in baseline activity within neural populations that encode effector-specific motor plans.

While these previous studies have provided evidence for limb repetition effects (same limb, different movement), but not movement repetition effects (same movement, different limb), an unresolved question is whether repetition of both aspects (i.e., limb and movement) yields an additional RT benefit. Such an effector-specific repetition effect of movement parameters would be of theoretical interest because it would imply a hierarchical organization of motor planning (Wong et al., 2015), that is, imply that movement plans can be recycled, but only by a given limb and not across limbs. Thus, the fact that limb repetition improves reaction times independent of the specific movement implies the existence of limb-specific resources; in contrast, effector-specific repetition advantages of a specific movement would suggest that movement parameters are linked to and specified after the effector has been chosen, and not coded in a body-independent manner (Seegelke et al., 2014; Swinnen et al., 2010; Wong et al., 2015).

Indeed, combined repetition of limb and movement have yielded by far the fastest RTs in the respective studies (Seegelke et al., 2021; Valyear & Frey, 2014). However, these studies have confounded repetition of motor and visual aspects: in the respective experiments, participants learned to associate a visual cue with each action; when prime and probe were repeated to cue repetition of both limb and movement, this implied also repeating the visual cue that had been associated with the respective action (Seegelke et al., 2021; Valyear & Frey, 2014). Therefore, it is possible that the large RT benefit of combined repetition was attributable to visual processing of the cue and/or associative rule retrieval, rather than to reuse of motor plans. This conjecture appears rather plausible given that visual priming/repetition effects are a well-known and robust phenomenon (Grill-Spector et al., 2006; Schacter & Buckner, 1998; Schacter et al., 2004). Accordingly, to obtain a pure readout of motor plan reuse, it is necessary to avoid visual repetition, an aspect not accounted for in previously reported studies.

To this end, we modified the paradigm of our previous study (Seegelke et al., 2021). Participants performed two movements in succession, a prime and a probe action. They used either the left or the right hand and moved it in one of two directions, inward or outward with respect to the body midline. The key design feature of the present experiment is that different visual cues were used for prime and probe actions. Thus, each of the four possible actions was associated with two specific visual symbols, one for prime and one for probe actions (Figure 1). Thus, all prime-probe combinations, including those involving a combined repeat of limb and movement, were cued with two different symbols; our manipulation, thus, removes the confound of visual repetition that was present in earlier studies. We previously obtained similar repetition effects for different effectors – hands and feet – as well as the phrasing of movement direction instructions as either egocentric (inward vs. outward) or allocentric (leftward vs. rightward). To keep working memory load due to associating stimuli and responses identical to our previous study which used identical symbols for prime and probe (Seegelke et al., 2021), we did not include foot movements in the present study, for which primes and probes were associated with different stimulus sets. Moreover, we restricted our design to the egocentric instruction to save acquisition time.

**Figure 1.**
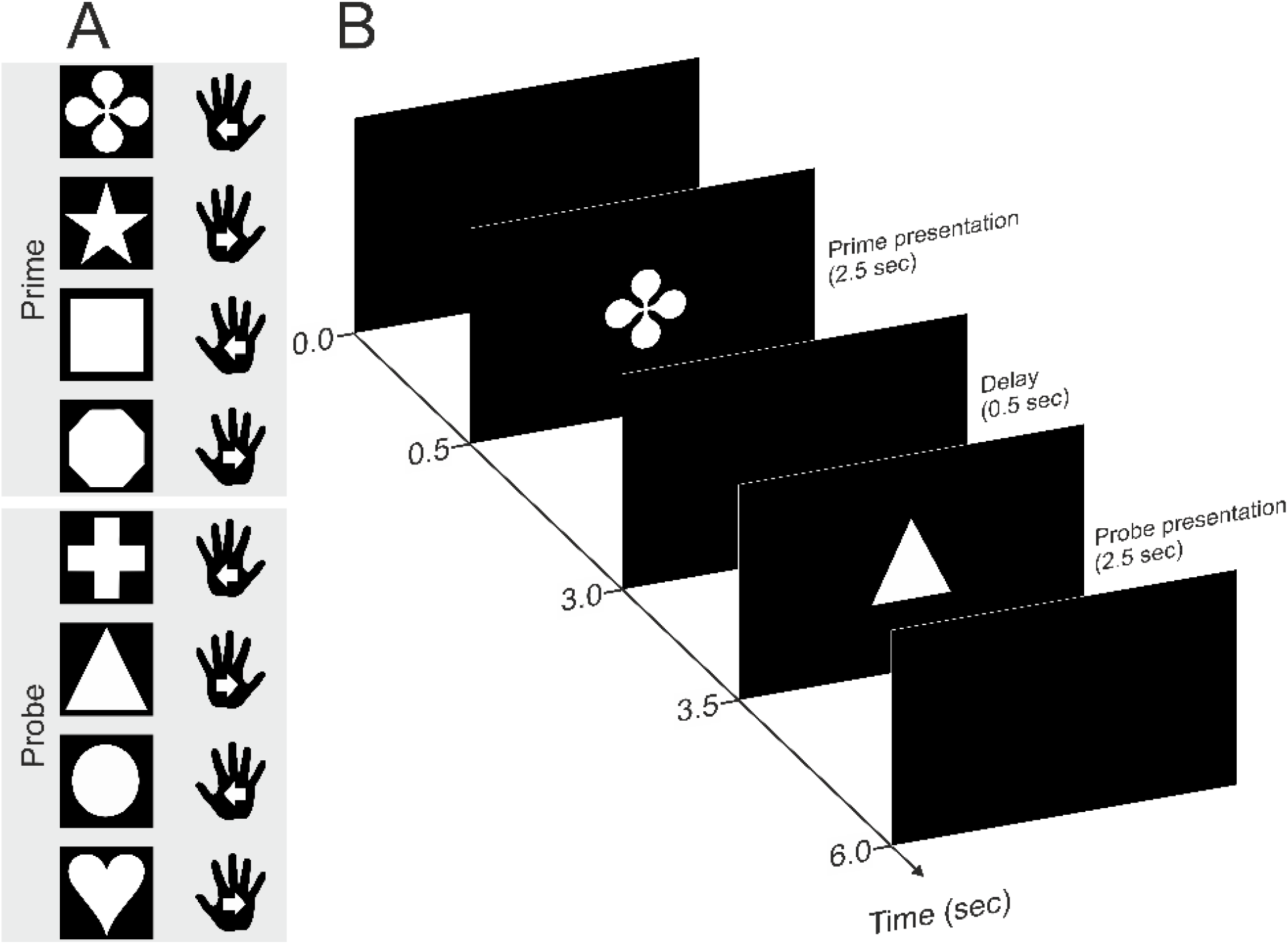
Experimental task and procedure. Participants performed two successive movements (a prime and a probe) separated by a 0.5-s delay interval. Movements were performed in one of two directions (inward, outward) with either hand (left, right). A: example shape-action assignment. Actions were defined through arbitrary rules by means of different shapes visually displayed on a computer screen. Each shape corresponded to two specific actions (one for the prime and one for the probe phase), and shape action associations were randomized across participants. White arrows indicate movement direction. B: example trial sequence with event timing.

Based on previous findings, successive, i.e., probe, actions should be initiated faster when participants use the same hand, even when the specific motor parameters, operationalized as movement direction, are different between prime and probe. More importantly, however, the idea of a processing hierarchy implies that participants should be fastest if both limb and movement are repeated.

## Methods

### Open Science and Ethics Statement

The study was preregistered on the Open Science Framework website (https://osf.io/rzt6j). Data and code for the present article are available at the Open Science Framework website https://osf.io/e9up2/. The University of Salzburg Ethics committee approved all procedures (Ethical Application Ref: EK-GZ 32/2023).

### Participants

We defined a target sample size of N = 20, because we obtained significant results with 20 participants in a previous study using similar procedures (Seegelke et al., 2021). However, results were statistically inconclusive with the pre-registered sample size (see Supplementary Information); therefore, we collected data from another 20 participants. Thus, our final sample comprised 40 students (self-reported gender: 27 female, 12 male, 1 non-binary, mean age = 24.18 years, SD = 9.60) which participated in exchange for course credit. 38 participants were right-handed (mean handedness score = 91.87, SD = 11.79), two participants were left-handed (mean handedness score = -60, SD = 56.57; Oldfield). All participants were physically and neurologically healthy and had normal or corrected-to-normal vision according to self-report.

### Apparatus and stimuli

Participants sat in front of a 24-in computer monitor (1920 × 1200 pixels; 60-Hz refresh rate) that was positioned on a table at a viewing distance of about 75 cm. Two square PVC blocks (10 × 10 × 3cm) with centrally embedded round pushbuttons (7cm in diameter) served as start location for the left and right hand. They were positioned about 40 cm apart (center-to-center distance) at the table’s front edge equidistant with respect to the body midline within comfortable distance.

Participants performed two successive movements (i.e. a prime and a probe action) in response to visual white shapes (413 × 413 pixels) presented centrally on the monitor against a black background (Figure 1). Each shape defined one of four responses: an inward left-hand movement, an outward left-hand movement, an inward right-hand movement, or an outward right-hand movement, with respect to the body midline. Two avoid stimulus repetition between prime and probe actions, each of the four actions was associated with two visual specific shapes, one for the shape as prime and one for the shape as probe), with the specific shape-action assignment randomized across participants. The experiment was controlled with MATLAB (version R2015; The MathWorks, Natick, MA) using the Psychophysics Toolbox (Brainard, 1997). An optical motion capture system (Visualeyez II VZ4000v; Phoenix Technologies Inc., Vancouver, BC, Canada) recorded hand kinematics at 250-Hz sampling rate. We placed infrared markers on the dorsal side of each hand (distal end of the third metacarpal).

### Procedure

At the start of each trial, participants depressed the start buttons with their hands for 500 ms. The prime stimulus, i.e., one of the four shapes for prime actions, then appeared on the screen for 2500 ms and participants performed the respective action and then moved the hand back to the start position. After a 500 ms delay, the probe stimulus (one of the other four shapes for probe actions) appeared and participants then executed the respective probe action. A 2000 ms inter-trial interval followed, after which participants could initiate the next trial by depressing the buttons. If participants did not complete the respective action within the 2500 ms stimulus display interval or released the wrong response button, they received the error message “Too slow” or “Wrong effector”, respectively, on the screen. We instructed participants to perform inward and outward movements (with respect to the body midline) of about 20 cm at a comfortable speed. On average, participants moved their hands 19.3 cm and 19.4 cm for prime and probe movements, respectively. Movement distances were also similar across hands and movement direction (see Supplemental Table S2). The 16 possible prime-probe combinations were repeated four times in each block in a randomized order, and each participants completed five blocks, yielding 320 trials in total.

Prior to the experimental blocks, participants completed one to two practice blocks until they reported to be familiar with the stimulus-response associations. During practice blocks, stimulus-response associations were visible to the participants. The procedure was identical to the main experiment except that participants performed only one action in response to a single stimulus. Each stimulus was repeated eight times within a practice block for a total of 64 trials. On average, participants performed the wrong action in 4.9% of practice trials. RT data over the course of the practice trials are shown in Supplemental Figure S3. It is evident that RTs similarly decreased for all responses and were about level (and equal for all responses) towards the end of practice trials The experiment took about 1.5 hours to complete.

### Data processing and analysis

We processed kinematic data with customized script in MATLAB (version R2021b; The MathWorks, Natick, MA). We interpolated missing data points using the spring metaphor method in the inpaint_nan function (D’Errico, 2021) and low-pass filtered the data using a second-order Butterworth filter with a cutoff frequency of 6 Hz. We determined movement onset as the time point at which the vectorial velocity of a marker exceeded 50 mm/s and reaction time (RT) as the time between stimulus onset and movement onset. We excluded trials in which no data were recorded and trials in which trajectories could not be reconstructed because too many data points were missing (286 trials, 2.2%). We removed trials in which participants performed the wrong prime or probe action or did not complete the respective action within the 2500 ms stimulus display interval (857 trials, 6.7%) and trials in which RT of probe actions was faster than 200 ms or slower than 1,500 ms (46 trials, 0.4%).

### Statistical approach

To analyze our data, we fitted Bayesian regression models created in Stan (http://mc-stan.org/) and accessed with the package brms version 2.16.3 (Bürkner, 2017) in R (R Development Core Team, 2020). In a first step, we fitted Bayesian regression models with RT as dependent variable and the within-subject variables Hand (left, right) and Movement Direction (inward, outward), separately for prime and probe actions, to assess whether RT differed for the different movement types. For our main analysis, we fitted a model with probe RT as dependent variable and the within-subject categorical variables Hand Repeat (repeat, switch) and Movement Direction Repeat (repeat, switch). We set orthogonal contrasts using the set_sum_contrast() command in afex version 1.0-1 (Singmann et al., 2020) and included random intercepts and slopes. We used a shifted log-normal distribution to estimate RTs and specified mildly informative priors for population-level (i.e., fixed) effects: For the intercept, the prior was a normal distribution with mean 6.4 and SD 0.5 (log-scale); for the effect of Hand Repeat, the prior was a normal distribution with mean 0.055 and SD 0.1 (log-scale). These settings reflect the prior knowledge of a main effect of hand repetition of about 70 ms, as we found in a previous study (Seegelke et al., 2021). To compare our results to findings of a previous study in which visual stimuli could be repeated between prime and probe (Seegelke et al., 2021), we additionally fitted a model with probe RT as dependent variable, the within-subject categorical variables Hand Repeat (repeat, switch) and Movement Direction Repeat (repeat, switch), and the between-subject categorical variable Experiment (Present Experiment, Seegelke et al., 2021 Experiment 1; Seegelke et al., 2021 Experiment 2). The estimation of parameters’ posterior distributions was obtained by Hamiltonian Monte-Carlo sampling with 4 chains, 1,000 sample warmup, and 11,000 iterations and checked visually for convergence (high ESS and Rhat ≈ 1). We used the package bayestestR version 0.11.5 (Makowski, Ben-Shachar, Chen, & Lüdecke, 2019; Makowski, Ben-Shachar, & Lüdecke, 2019) to describe the parameters of our models. We report the median as a point estimate of centrality and the 95% credible interval (CI) computed based on the highest-density interval (HDI) to characterize the uncertainty related to the estimation. As an index of existence of an effect, we report the Probability of Direction (pd), representing the certainty associated with the most probable direction (positive or negative) of the effect (Makowski, Ben-Shachar, Chen, & Lüdecke, 2019). Following the recommendations of Makowski, Ben-Shachar, Chen, and Lüdecke (2019), for interpretation we consider 95%. 97%, and 99% as reference points for the pd (pd <= 95%: uncertain; pd > 97%: likely existing; > 99%: probably existing). In addition, as an index of the significance of a given effect, we tested whether the HDI excluded a region of practical equivalence (ROPE) of ±0.1 effect sizes around 0. If the HDI is completely outside the ROPE, the null hypothesis for this effect is rejected. If the ROPE completely covers the HDI (i.e., all most credible values of a parameter are inside the ROPE), the null hypothesis is accepted.

## Results

Table 1 shows median RTs for the different movement types (i.e., left and right hand, inward and outward movement direction) for both prime and probe actions. Bayesian analysis confirmed that RT was similar for the left and right hand (Prime movement: median difference = 12 ms, 95% HDI [-10,33], 66% of HDI in ROPE, pd = 87.02%; Probe movement: median difference = -1 ms, 95% HDI [-22,19], 89% of HDI in ROPE, pd = 55.06%), as well as for inward and outward movements (Prime movement: median difference = 20 ms, 95% HDI [-4,43], 48% of HDI in ROPE, pd = 95.55%; Probe movement: median difference = 14 ms, 95% HDI [-4,31], 64% of HDI in ROPE, pd = 94.02%). These results indicate that RT was similar independent of the experiment’s the parameters.

**Table 1.** RTs (in ms) for prime and probe actions as a function of hand and movement direction. Estimated median values and 95% highest density interval (HDI).

| **Prime action** |  |  |  |
| --- | --- | --- | --- |
| Hand | Movement Direction | Median | 95% HDI |
| Left | Inward | 665 | 629 – 701 |
| Left | Outward | 637 | 605 – 671 |
| Right | Inward | 645 | 612 – 681 |
| Right | Outward | 632 | 600 - 666 |
| **Probe action** |  |  |  |
| Left | Inward | 655 | 622 – 688 |
| Left | Outward | 640 | 608 – 672 |
| Right | Inward | 655 | 623 – 689 |
| Right | Outward | 642 | 610 - 675 |

Figure 2 shows median RTs for probe actions as a function of Hand Repeat and Movement Direction Repeat. To assess potential differences statistically, we calculated contrasts between the posterior distributions of the respective conditions. In general, RTs were shorter for Hand Repeat trials than for Hand Switch trials (median difference = 55 ms, 95% HDI [46,63]; 0% of HDI in ROPE, pd = 100%), indicative of a limb repetition effect. The limb repetition effect was statistically reliable when calculated separately for Movement Repeat trials (median difference = 73 ms, 95% HDI [60,87]; 0% of HDI in ROPE, pd = 100%) and Movement Switch trials (median difference = 36 ms, 95% HDI [26,46]; 0% of HDI in ROPE, pd = 100%, Figure 3A). RTs were also generally shorter for Movement Repeat trials than for Movement Switch trials (median difference = 26 ms, 95% HDI [19,35]; 0% of HDI in ROPE, pd = 100%), indicative of a movement repetition effect). Importantly though, these differences were statistically reliable only for Hand Repeat trials (median difference = 45 ms, 95% HDI [33,58]; 0% of HDI in ROPE, pd = 100%), but not for Hand Switch trials. For the latter comparison, we obtained strong support for the null hypothesis (median difference = 8 ms, 95% HDI [-2,17]; 100% of HDI in ROPE; pd = 94.09%, Figure 3B). Direct comparison demonstrated that the RT difference (Movement Direction Switch – Movement Direction Repeat) was significantly larger for Hand Repeat trials than Hand Switch trials (median difference = 37 ms, 95% HDI [22,54]; 0% of HDI in ROPE; pd = 100%). Together, these results replicate previous findings (Seegelke et al., 2021; e.g., Valyear & Frey, 2014) that successive movements are initiated faster when the same limb is used, even if a different movement is performed (i.e., limb repetition effect). Moreover, they show that repeating specific movement characteristics (e.g., movement direction) leads to an additional decrease in response latencies, but only if the movement is executed with the same limb as before.

**Figure 2.**
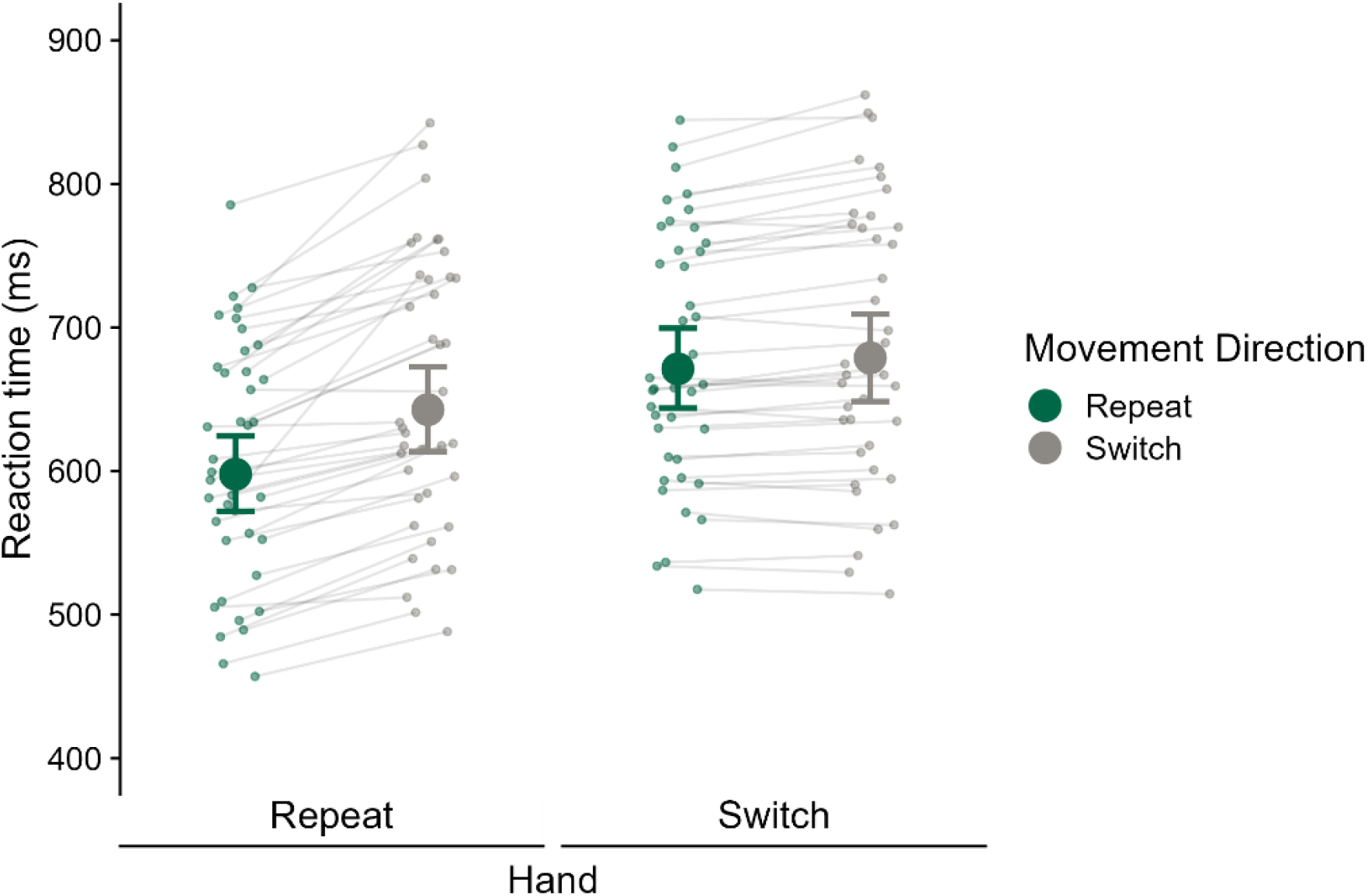
RTs (ms) for probe actions as a function of Hand and Movement Direction Repeat. Large and small dots represent estimated group and individual medians, respectively. Error bars reflect the 95% highest-density interval (HDI).

**Figure 3.**
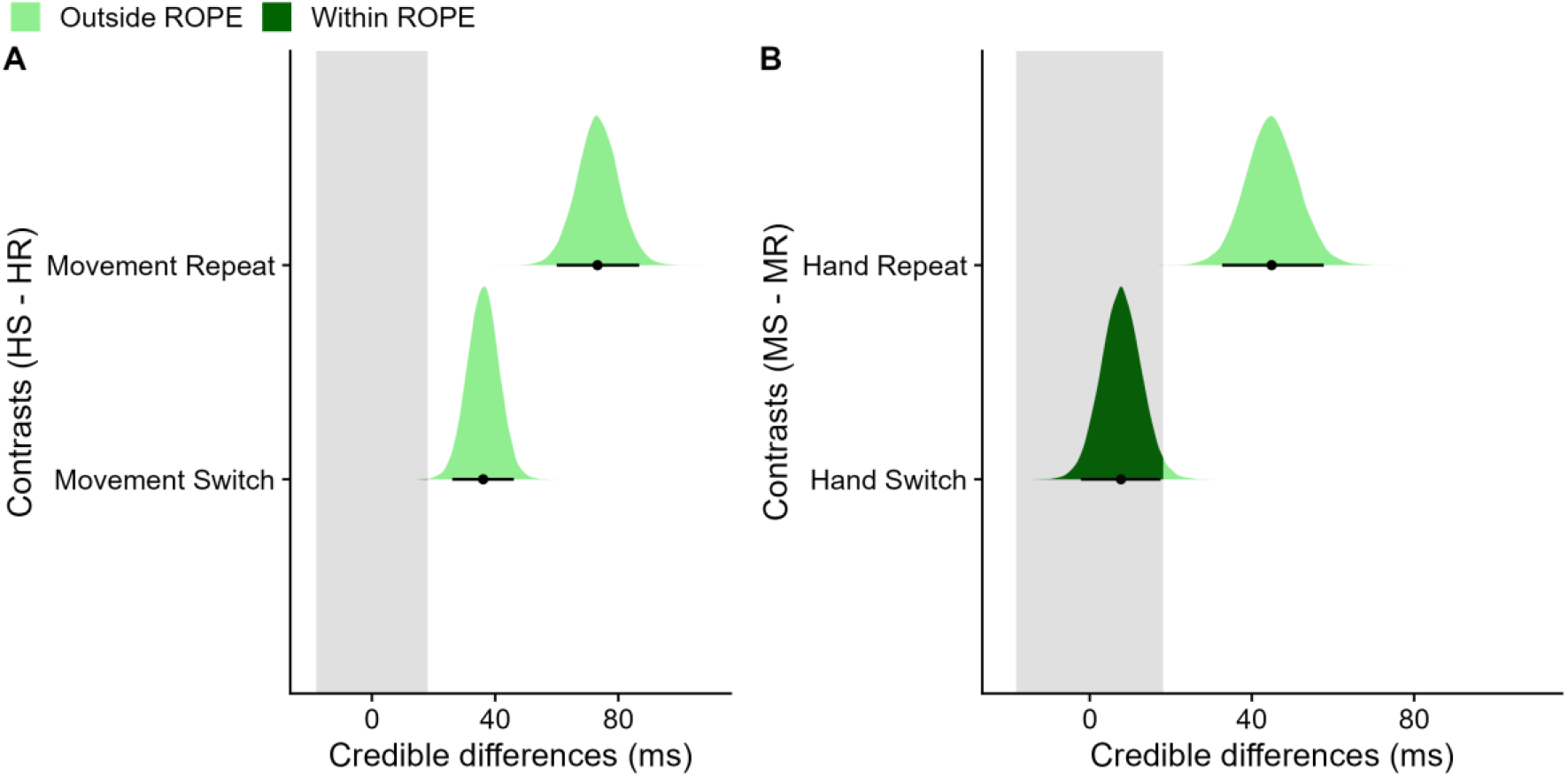
Assessment of limb repetition effect and movement repetition effect. Repeating the hand yielded reduced RTs regardless of whether the movement was repeated or switched. Repeating the movement yielded reduced RTs only when the same hand was used. A Contrasts of posterior distributions between trials with hand switch (HS) and trials with hand repetition (HR), separately for the movement direction repeat condition (top row) and the movement direction switch condition (bottom row). B Contrasts of posterior distributions between trials with movement direction switch (MS) and trials with movement direction repetition (MR), separately for the hand repeat condition (top row) and the hand switch condition (bottom row). Black dots represent the medians and error bars the 95% highest-density interval (HDI). The gray shaded area indicates the region of practical equivalence (ROPE) of ±0.1 effect sizes around 0 [-18, 18]. Portions of the distributions outside and inside the ROPE are shown in light green and dark green, respectively.

By reviewer request, we also analyzed the data with frequentist statistics using a 2 Hand (Repeat, Switch) × 2 Movement Direction (Repeat, Switch) repeated measures ANOVA. These analyses led to qualitatively similar conclusions as our Bayesian analysis (see Supplemental Table S3).

## Discussion

Previous work has shown that when two actions are performed in succession, response times to initiate the second action are faster when the same effector is used, even if movement parameters (e.g., grip type, movement direction) differ for the two actions (Seegelke et al., 2021; i.e., limb repetition effect; Valyear & Frey, 2014). However, a limitation of these studies was that they confounded motor and visual aspects of repetition because identical actions pairs were always instructed with identical visual cues. In the present study, we eliminated this confound by using non-identical visual cues to instruct identical action pairs, thus eliminating visual repetition and retaining only movement repetition in the experimental paradigm. We found that the limb repetition effect persisted when visual cue repetition was eliminated, that is, successive movements were initiated faster when the same limb was used whether or not movements were performed in the same direction. In addition, repeating movement direction also led to a decrease in response latencies, but only when the same hand was used for both actions.

While we did observe a combined hand and movement repetition effect, its size was considerably smaller than that reported previously (see Supplementary Figure S4 and S5 for a direct comparison to the results of Seegelke et al., 2021). Thus, while our study validates a motor origin of previously reported repetition effects, it nonetheless indicates that previous results were affected by visual repetition. Therefore, it is important to control for visual repetition to isolate motor-related effects, which can easily be done with the method we have introduced here.

Our findings complement previous work on the influence of the recent motor history on current motor performance. Motor history effects have been observed in different tasks and manifest as biases in hand choice (Schütz & Schack, 2020; Schweighofer et al., 2015; Valyear et al., 2019), movement characteristics such as grasp orientation (Kent et al., 2009; Schütz et al., 2011), reach direction (Diedrichsen et al., 2010; Jax & Rosenbaum, 2007; Tsay et al., 2022; Verstynen & Sabes, 2011), movement speed (Hammerbeck et al., 2014), and response latencies (Mawase et al., 2018; Seegelke et al., 2021; Valyear & Frey, 2014, 2015; Wong et al., 2017). According to the plan reuse hypothesis (Jax & Rosenbaum, 2007; Rosenbaum et al., 2007), these effects result from prior planning-related activity such that parameters of previously generated action plans are reused for current actions. This view is corroborated by studies that observed reduced response latencies of successive actions executed with the same hand (Seegelke et al., 2021; Valyear & Frey, 2014, 2015), or foot (Seegelke et al., 2021), and regardless of movement direction instruction (Seegelke et al., 2021), i.e., egocentric (inward vs. outward) vs. allocentric (leftward vs. rightward). Together, these findings have been taken to support the notion that repetition effects in action planning reflect encoding of effector-specific action plans.

In the present experiment, we observed a hand repetition effect for both identical and non-identical actions. Thus, the fact alone that the hand is used twice reduces RT, supporting the idea that action plans are effector-specific. Yet, a movement repetition effect was also evident, albeit contingent on both actions being performed with the same hand. Thus, the movement was apparently not coded independently but yoked to the hand that performed it. This pattern is in line with a hierarchical framework of motor planning (Wong et al., 2015) in which effector selection is distinct from, and precedes, movement specification. That is, participants first choose which hand to perform the movement with before they specify the motor parameters of the movement with the chosen hand. A similar hierarchy has been proposed for the processes of effector and target selection (Herbort & Rosenbaum, 2014). In this study, participants were free to reach to either of two targets with either hand. Performance in this free condition was more similar to a condition in which the hand was specified than to a condition in which the target was specified, supporting that effector selection preceded target selection. The conjecture of effector selection and movement specification being distinct processes is also consistent with brain imaging findings that place the two functions in different brain regions, namely effector selection in anterior regions of PPC (Leoné et al., 2014; Wong et al., 2015) and movement specification in premotor and primary motor regions (cf. Wong et al., 2015).

There is also evidence that certain movement features prone to the recent motor history can transfer from one hand to the other (Dixon et al., 2012; van der Wel et al., 2007). For example, in one study participants sequentially contacted several targets arranged in a semi-circle. On some trials, they had to move over an obstacle positioned between two targets. After obstacle clearance, participants’ peak movement heights only gradually returned to baseline, even when participants changed their hand after having moved over the obstacle (van der Wel et al., 2007). This finding has been interpreted as indicating the involvement of effector-independent representations in repetition effects and seems to contradict the present as well as previous studies that did not observe cross-hand transfer of motor parameters (Seegelke et al., 2021; Valyear & Frey, 2014, 2015).

We can only speculate about how these divergent findings can be reconciled. It has been suggested that optional motor planning process are required to resolve this ambiguity in situations in which the task does not constrain in which way the motor goal is achieved, (Wong et al., 2015). For example, some movement contexts might involve an abstract kinematic representation, in which the movement trajectory is planned in an effector-independent manner in extrinsic space. The existence of such an additional planning process is supported by numerous behavioral and neurophysiological studies across different motor tasks such has handwriting (Castiello & Stelmach, 1993; Rijntjes et al., 1999; Wing, 2000), drawing (Albert & Ivry, 2009), and reaching (Cluff & Scott, 2015; Wong et al., 2016). Thus, participants in the study by van der Wel (2007) may have formed an effector-independent representation of a desired trajectory over obstacles that was accessed by both hands, whereas movements in our study resembled point-to-point reaching movements that do not require specification of the abstract kinematics of a movement trajectory (Cluff & Scott, 2015; Wong et al., 2016; Wong et al., 2015). However, arguing against this possibility, there is also evidence for the involvement of effector-independent motor representations in movement such as key presses (Bapi et al., 2000), point-to-point reaching (Kumar et al., 2020), and simple flexing movements (Striem-Amit et al., 2018). Moreover, participants in the van der Wel study knew in advance that (and when) they had to change hands, whereas in our study the information which hand to use was given with the imperative stimulus. Thus, it is thinkable that the (RT-consuming) process of effector selection (Wong et al., 2015) masked manifestation of effector-independent representations in response latencies.

In conclusion we have eliminated an experimental confound that forbade strong interpretation of some previous results in previous studies of action repetition. We have shown that both a hand and a movement repetition effect are observable also after removal of this confound, which supports the proposition that motor planning integrates the recent motor history. Moreover, our experimental result of a hand-specific movement repetition effect suggests a hierarchical organization in which effector selection precedes movement specification.

## Acknowledgments

We thank Manuel Diaz, Esther Kraus, Carolin Mack, and Alica Mindarova-Hechl for help with data acquisition.

## Statements and Declarations

No conflicts of interest, financial or otherwise, are declared by the author(s).

## Author contributions

C. S. conceived and designed research; C. S. performed experiments; C. S. analyzed data; C. S. interpreted results of experiments; C. S. prepared figures; C. S. drafted manuscript; C. S. and T.H. edited and revised manuscript; C. S. and T.H. approved final version of manuscript.

