## Supplemental Material for "Effector selection precedes movement specification: evidence from repetition effects in motor planning"

### Supplementary Information

Here we report the results with our preregistered sample size of N = 20.

#### Participants

Participants (15 female, 5 male, mean age = 21.9, SD = 2.00) were right-handed (mean handedness score = 95.06, SD = 8.79; Oldfield, 1971), physically and neurologically healthy, and reported normal or corrected-to-normal vision.

#### Data processing and analysis

We excluded trials in which no data were recorded and trials in which trajectories could not be reconstructed because of too many missing data points (159 trials, 2.5%). We removed trials in which participants performed the wrong prime or probe action or did not complete the respective action within the 2500 ms stimulus display interval (362 trials, 5.7%) and trials in which RT of probe actions was faster than 200 ms or slower than 1,500 ms (23 trials, 0.4%).

#### Results

RT was very similar across the different movement types (i.e., left and right hand, inward and outward movement direction) for both prime and probe actions (Table S1). Bayesian analysis with Hand (left, right) and Movement direction (inward, outward) as within-subject predictors confirmed that the HDI was inside the ROPE for all predictors for both prime and probe actions (Figure S1). This result demonstrates that none of the parameters affected RT.

**Table S1.** RTs (in ms) for prime and probe actions as a function of hand and movement direction. Estimated median values and 95% highest density interval (HDI).

| **Prime action** | | | |
|---|---|---|---|
| Hand | Movement Direction | Median | 95% HDI |
| Left | Inward | 664 | 617 – 711 |
| Left | Outward | 647 | 612 – 685 |
| Right | Inward | 647 | 597 – 699 |
| Right | Outward | 645 | 602 - 691 |
| **Probe action** | | | |
| Left | Inward | 648 | 604 – 698 |
| Left | Outward | 631 | 588 – 674 |
| Right | Inward | 640 | 598 – 686 |
| Right | Outward | 638 | 592 - 687 |

Figure S1 shows median RTs for probe actions as a function of hand repeat and movement direction repeat. RTs were shorter when the hand was repeated (median = 614 ms, 95% HDI [577,653] than when it was not repeated (median = 666 ms, 95% HDI [627,708]). RTs were also shorter when successive actions comprised the same movement direction (median = 627 ms, 95% HDI [591,664] rather than different movement directions (median = 653 ms, 95% HDI [612,696]). Importantly, during hand repeat trials, RTs were considerable shorter when movement direction was also repeated (median = 593 ms, 95% HDI [558,631] compared to when movement direction was not repeated (median = 634 ms, 95% HDI [593,675]). In contrast, during trials in which hand was not repeated, the

RT difference between trials in which movement direction was repeated (median = 660 ms, 95% HDI [621,699] and trials in which it was not repeated (median = 672 ms, 95% HDI [630,717]) was much smaller. We calculated contrasts between the posterior distributions trials in which movement direction was repeated and trials in which it was not repeated, separately for hand repeat and hand no repeat trials (Figure S2). For hand repeat trials, there was a substantial RT difference (median difference = 41 ms, 95% HDI [21,61]; 0% of HDI in ROPE, pd = 99.99%). For hand no repeat trials, the RT differences was much smaller (median difference = 12 ms, 95% HDI [-4,28]; 67% of HDI in ROPE; pd = 92.92%). However, although direct comparison revealed that the RT differences are larger in hand repeat than the RT differences in hand no repeat trials, a considerable portion of the HDI was still inside the ROPE and the pd indicated that the effect is likely existing (median difference = 29 ms, 95% HDI [3,55]; 28% of HDI in ROPE; pd = 98.56%).

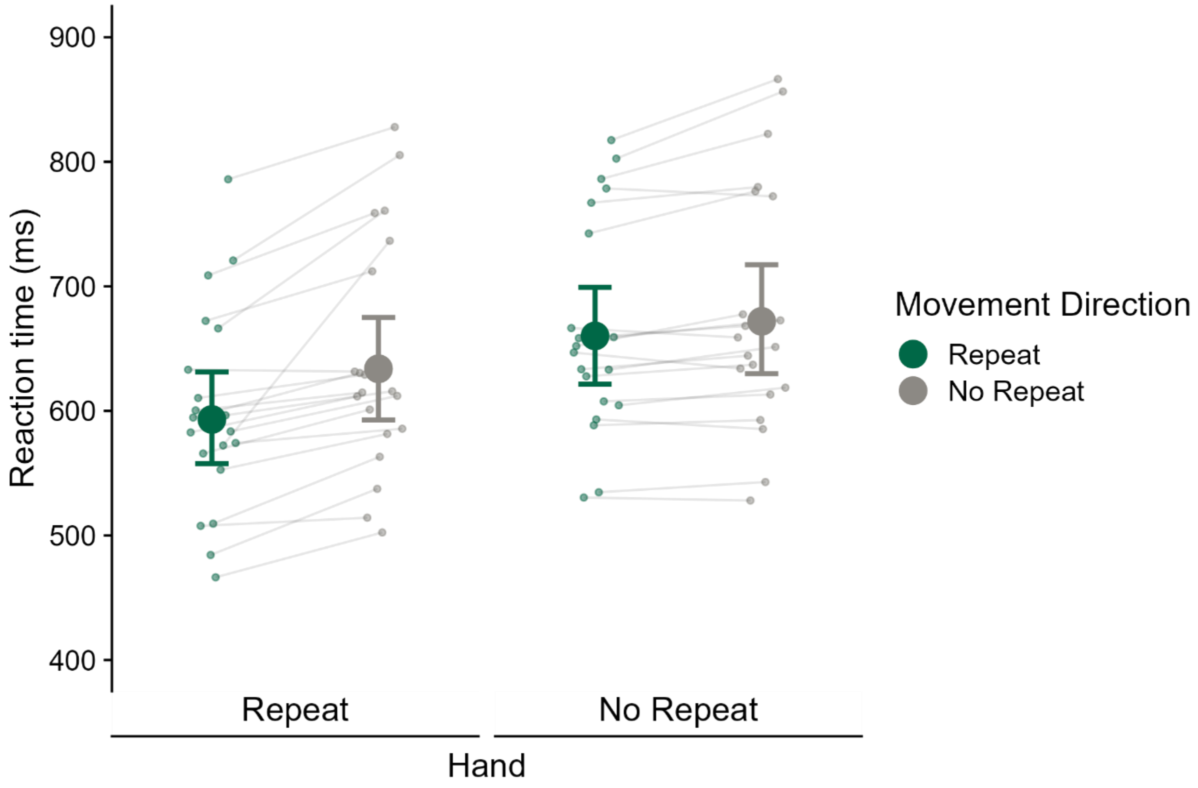


**Figure S1.** RTs (ms) for probe actions as a function of hand and movement direction repeat based on our preregistered sample size (N = 20). Large and small dots represent estimated group and individual medians, respectively. Error bars reflect 95% highest-density interval (HDI).

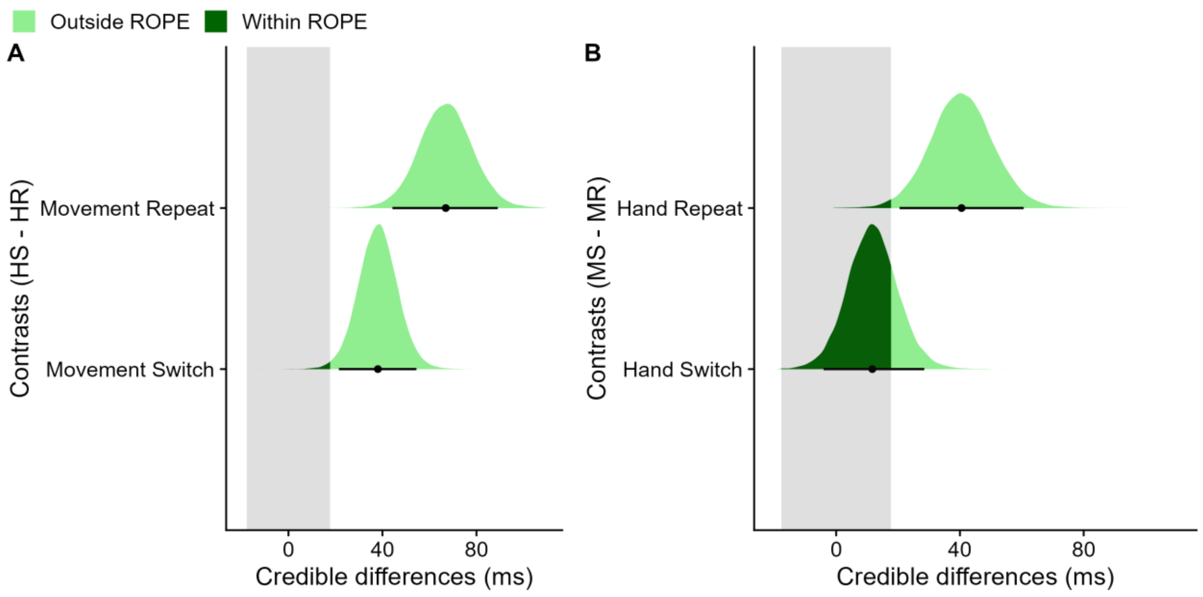


**Figure S2.** Assessment of limb repetition effect and movement repetition effect based on our preregistered sample size (N = 20). Repeating the hand yielded reduced RTs regardless of whether the movement was repeated or switched. Repeating the movement yielded a larger reduction in RTs when the hand was repeated than when it was switched (median difference = 29 ms). However, direct comparison demonstrated that the RT difference (Movement Direction Switch – Movement Direction Repeat) was statistically not reliable (95% HDI [3,55]; 28% of HDI in ROPE; pd = 98.56%). **A** Contrasts of posterior distributions between trials with hand switch (HS) and trials with hand repetition (HR), separately for the movement direction repeat condition (top row) and the movement direction switch condition (bottom row). **B** Contrasts of posterior distributions between trials with movement direction switch (MS) and trials with movement direction repetition (MR), separately for the hand repeat condition (top row) and the hand switch condition (bottom row). Black dots represent the medians and error bars the 95% highest-density interval (HDI). The gray shaded area indicates the region of practical equivalence (ROPE) of ±0.1 effect sizes around 0 [-18, 18]. Portions of the distributions outside and inside the ROPE are shown in light green and dark green, respectively.

The following results are based on our final sample size (N = 40).

**Table S2.** Movement distances (in cm) of prime and probe actions measured in x-direction (n = 40 participants).

| **Prime action** | | | |
|---|---|---|---|
| **Hand** | **Movement Direction** | **M** | **SD** |
| **Left** | **Inward** | **18.2** | **4.6** |
| **Left** | **Outward** | **20.8** | **5.5** |
| **Right** | **Inward** | **18.7** | **4.5** |
| **Right** | **Outward** | **19.7** | **5.7** |
| **Probe action** | | | |
| **Left** | **Inward** | **18.2** | **4.6** |
| **Left** | **Outward** | **21.0** | **5.4** |
| **Right** | **Inward** | **18.7** | **4.5** |
| **Right** | **Outward** | **19.6** | **5.9** |

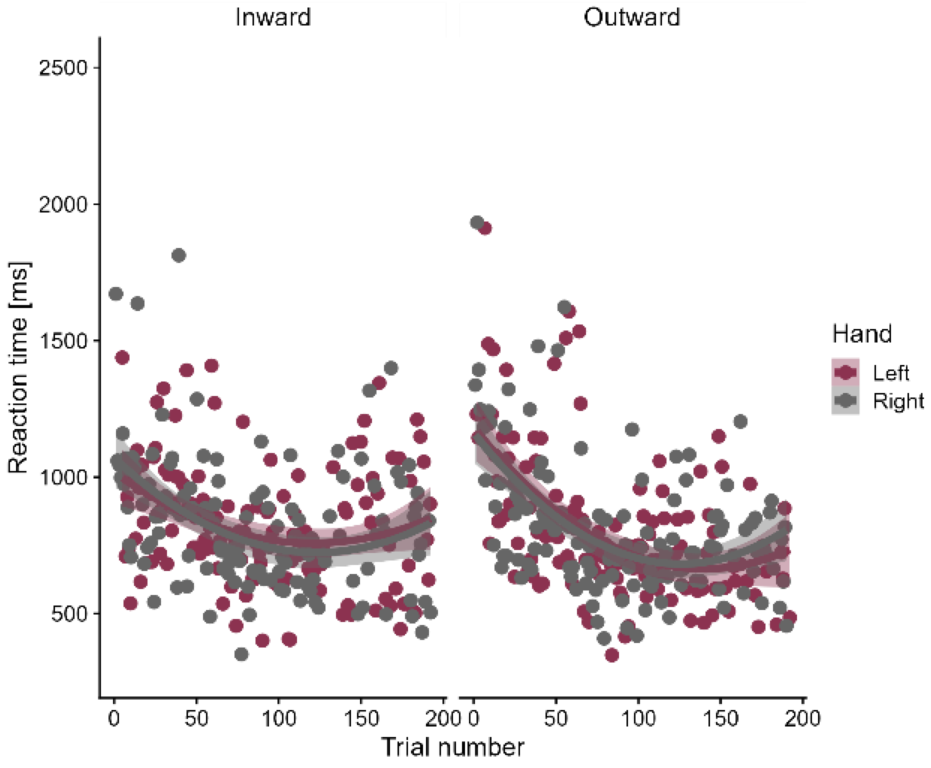


**Figure S3.** RTs (in ms) during practice trials as a function of Hand and Movement Direction showing that RT decreases across practice trials. Dots represent the trial medians; the lines show quadratic fits to the data.

**Table S3.** Outcome of the 2 Hand (Repeat, Switch) x 2 Movement Direction (Repeat, Switch) RM ANOVA with median RT of probe action (n = 40 participants). Analysis revealed significant main effects of both factors and their interaction. Similar to our Bayesian contrast analysis, follow-up simple effects tests showed that the limb repetition effect (i.e., Hand Switch minus Hand Repeat) was statistically significant when calculated separately for Movement Repeat trials (median difference = 60 ms, p <.0001) and Movement Switch trials (median difference = 33 ms, p <.0001). In contrast, the movement repetition effect (i.e., Movement Direction Switch – Movement Direction Repeat) was statistically significant only for Hand Repeat trials (median difference = 36 ms, p <.0001) but not Hand Switch trials (median difference = 9 ms, p =.0682). Df = degrees of freedom, MSE =mean squared error.

| **Effect** | **Df** | **MSE** | **F-value** | **Partial eta²** | **p-value** |
|---|---|---|---|---|---|
| Hand (H) | 1,39 | 607.11 | 140.44 | .783 | <.001 |
| Movement direction (M) | 1,39 | 890.06 | 22.71 | .368 | <.001 |
| H × M | 1,39 | 578.30 | 12.29 | .240 | <.001 |

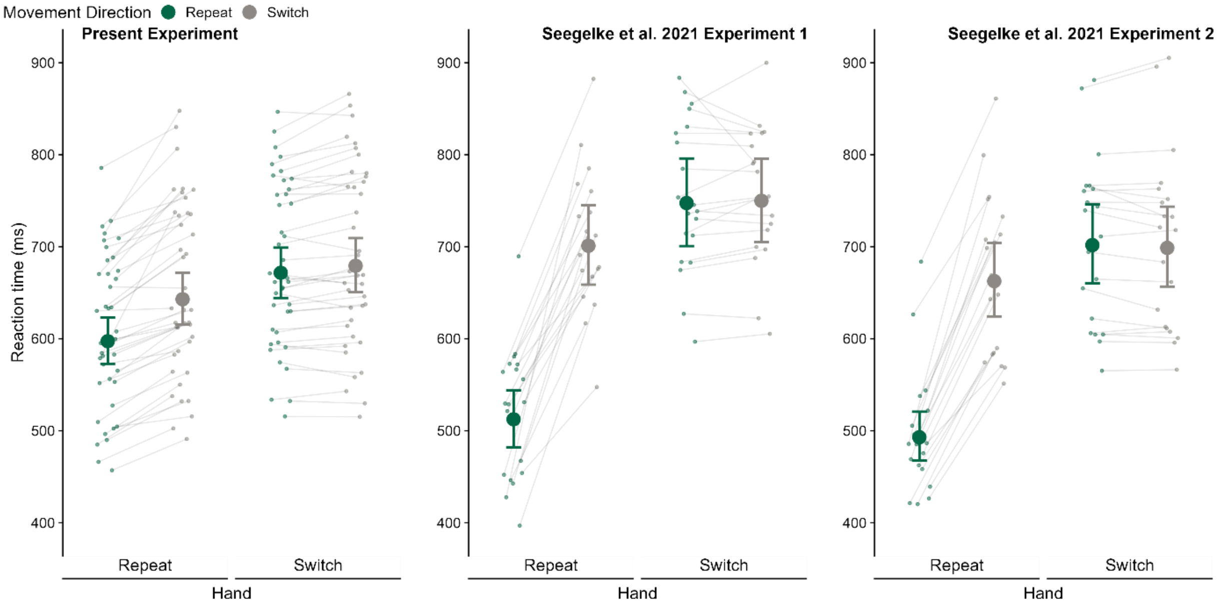


**Figure S4.** RTs (ms) for probe actions as a function of Hand and Movement Direction Repeat for the present experiment (left panel) and experiment 1 (middle panel) and experiment 2 (right panel) of Seegelke et al., (2021). Large and small dots represent estimated group and individual medians, respectively. Error bars reflect the 95% highest-density interval (HDI).

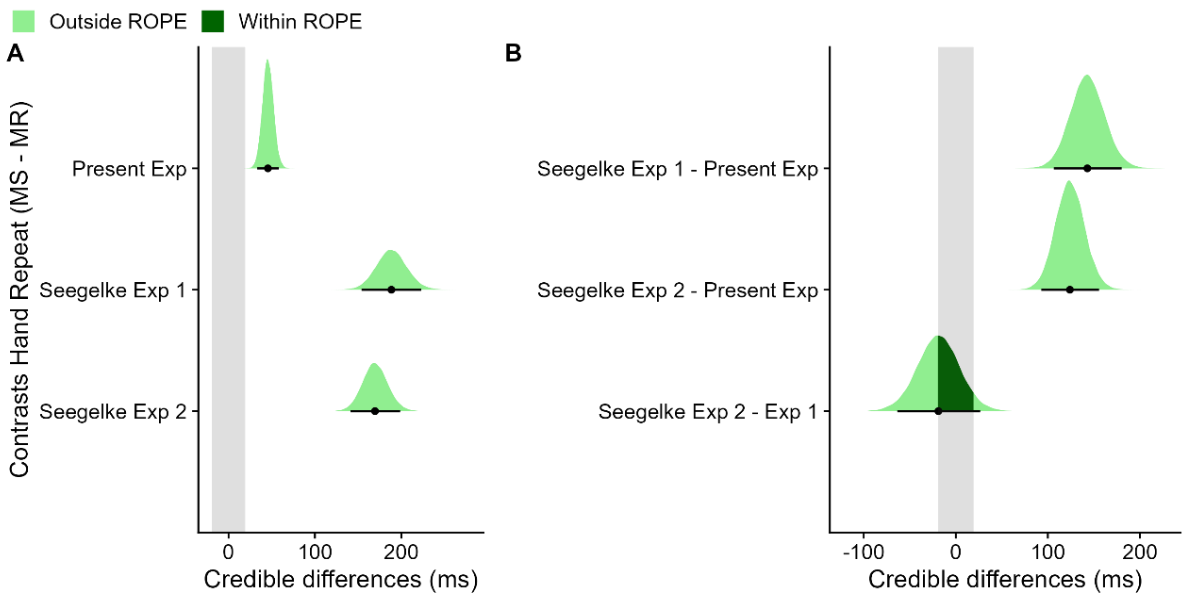


**Figure S5.** Comparisons of combined hand and movement repetition effect between the present experiment and experiment 1 and experiment 2 of Seegelke et al., (2021). The size of the effect in the present study, which controlled for repetition of visual cues, was considerably smaller than the effects in Seegelke et al., (2021), in which repetition of both hand and movement direction also comprised visual cue repetition. **A** Contrasts of posterior distributions between trials with movement direction switch (MS) and trials with movement direction repetition (MR), for the hand repeat conditions in the present experiment (top row: median difference = 46 ms, 95% HDI [33,58]; 0% of HDI in ROPE; pd = 100%), experiment 1 (middle row: median difference = 189 ms, 95% HDI [154,223]; 0% of HDI in ROPE; pd = 100%), and experiment 2 (bottom row: median difference = 170 ms, 95% HDI [141,199]; 0% of HDI in ROPE; pd = 100%) of Seegelke et al., (2021). **B** Contrasts of posterior distributions between the effects in panel A. Seegelke et al. 2021 Experiment 1 vs. Present experiment (top row: median difference = 143 ms, 95% HDI [106,180]; 0% of HDI in ROPE; pd = 100%); Seegelke et al. 2021 Experiment 2 vs. Present experiment (middle row: median difference = 124 ms, 95% HDI [93,156]; 0% of HDI in ROPE; pd = 100%); Seegelke et al. 2021 Experiment 2 vs. Seegelke et al. 2021 Experiment 1 (bottom row: median difference = -19 ms, 95% HDI [-64,26]; 43% of HDI in ROPE; pd = 80%). Black dots represent the medians and error bars the 95% highest-density interval (HDI). The gray shaded area indicates the region of practical equivalence (ROPE) of ±0.1 effect sizes around 0 [-19, 19]. Portions of the distributions outside and inside the ROPE are shown in light green and dark green, respectively.
